# State-Dependent Organization of Microscale Functional Circuitry in Visual Cortex

**DOI:** 10.64898/2026.03.31.715474

**Authors:** Rahul Biswas, Hasika Wickrama Senevirathne, Yujing Wang, Jiang Zhang, Somabha Mukherjee, Reza Abbasi-Asl

## Abstract

Directed functional connections between individual neurons, inferred from recorded activity, reflect information routing in cortex. Yet their organization across brain states and relation to synaptic connectivity remain poorly understood. Here, we present one of the largest single-neuron-resolution directed functional circuit maps, across about 45,000 neurons in four mouse visual areas. We inferred connections from deconvolved two-photon calcium activity with CITS, a causal-inference method designed for neural time series. In co-registered electron microscopy, CITS-inferred pairs were 3.0-fold more likely to be synaptically connected than other pairs. The map’s organization aligned with synaptic connectivity at areal, laminar, and cell-type levels. In low and high pupil-linked arousal, the anterolateral area (AL) had the highest intra-areal density. The strongest inter-areal pathway linked AL with the rostrolateral area. This organization held in both states, yet mean connection strength was higher in high arousal in every scan, by 9.4% on average. Finally, stronger connections linked neurons with more similar stimulus-driven predictive performance in both states.

## 1 Introduction

Microscale *functional circuitry*, defined here as directed and statistically causal functional connections between individual neurons inferred from neuronal activity, provides a window into the brain’s circuit-level organization [1,2]. A large body of theoretical and computational studies suggests that functional circuitry captures the dynamic routing of information through the brain [3–7]. Traditional functional connectivity methods have relied on undirected associations and cannot capture directional interactions [8]. Recent methods utilize statistical causal inference frameworks for neural activity recordings, enabling the estimation of directed functional connections and advancing the field from associative connectivity toward microscale functional circuitry [8–11]. Despite this progress, it remains unknown how such directed functional circuitry is organized across different brain states, particularly at the resolution of individual neurons.

Neurons’ functional interactions align only partially with their synaptic wiring. In *C. elegans*, whose synaptic connectome is fully reconstructed, a directed map of signal propagation obtained by activating single neurons and recording the responses of the rest departs from its synaptic connectome in both directions. Some synaptically wired pairs carry no signal, while some strongly responding pairs are linked only by extrasynaptic neuropeptide signaling [12]. Such exhaustive neuron-by-neuron mapping does not scale beyond small populations, so in mammalian cortex directed interactions are inferred from recorded activity. In mouse visual cortex, structure-function studies have so far related undirected response similarity to synaptic connectivity, showing neurons with correlated responses and similar tuning are preferentially and more strongly synaptically connected [13, 14]. Directed connectivity has been inferred from regional fMRI and field-potential signals, and from spike trains in small neural populations [8, 15–17], but not at single-neuron resolution across a dense connectome. As a result, the organization of these directed functional connections at single-neuron scale, and their relationships to anatomical connectivity, have not been examined systematically across cell types, layers, and visual areas.

Arousal state is a particularly important dimension along which to examine this question. Neural responses during sensory processing, attention, and behavioral tasks are known to vary with arousal, ranging from drowsiness to alertness [18–20]. Prior studies that examined functional connectivity across brain states relied on local field potential (LFP) array signals [21] or combined multi-unit spiking and LFP recordings [22]. While these revealed the state-dependent nature of population-level coordination, neither approach could resolve how circuits are organized at the level of individual neurons. Therefore, how directed functional circuitry is shaped across arousal states, at the level of individual neurons, cell types, laminar origin, and multi-area hierarchy, remains an open question.

To answer these questions, we leveraged the MICrONS dataset [14, 23], a unique resource combining large-scale calcium imaging of 146,388 neurons, of which 15,434 were co-registered with a dense electron microscopy (EM) reconstruction. This allowed us to map the functional circuitry of the mouse visual cortex with single-neuron resolution, estimating state-dependent connections across four visual areas (primary visual cortex V1, lateromedial LM, anterolateral AL, rostrolateral RL visual areas) and linking them to their anatomical positions, structural scaffold [24, 25], and cell types [26]. To infer directed functional connections from the neural activity recordings, we employed CITS (Causal Inference in Time Series) [9, 27], a statistical causal-inference method that is specifically designed for neural time series. Pupil size is a proxy for internal arousal, tracking noradrenergic and cholinergic activity [28, 29]. We defined arousal states by a pupil index, the part of pupil size not explained by the visual stimulus (luminance, contrast, motion and stimulus type). Within each scan, we labeled each imaging frame (about 159 ms) high-arousal if the index was at or above the scan median, and low-arousal otherwise. We kept only stretches of one state lasting at least 10 s (Methods). We mapped the directed *statistically causal functional circuitry* (CFC) within each arousal state across visual areas, cell types, and layers.

The directed functional connections between neuron pairs were 3.0-fold more likely to be synaptically connected than absent connections, in both arousal states. The functional circuit itself was intra-areal dominant. Intra-areal connections were denser than inter-areal connections in both arousal states, with area AL showing the highest within-area density and the AL*↔*RL axis as the prominent inter-area pathway. Across layers, within-area connections were densest in layer 5. At the areal, laminar and cell-type levels, these emergent functional patterns matched the organization of the co-registered synaptic connectome. Both arousal states showed this organization but differed in connection strength. Functional connection strength was higher in high-arousal than in low-arousal periods in all 12 scans and all five sessions, by 9.4% on average. The increase was significant in 12 of 16 directed area pairs and in every layer within areas. For stimulus-driven response prediction, neuron pairs with stronger functional connections exhibited more similar predictive performance in both states, with performance varying by cortical layer and cell type.

## 2 Results

### 2.1 CITS-inferred microscale functional circuitry is aligned with the synaptic connectome

The architecture of cortical functional circuitry, i.e., which pathways are active and their strength, shapes how sensory information is routed within and across visual areas [24, 30]. Cortical circuits are organized around dense local connectivity and sparser long-range connections [24, 31]. We first establish the baseline architecture of directed functional circuitry in each arousal state, and ask how this architecture varies by area and whether it is consistent across arousal states. Arousal state was continuously monitored via pupil size [18, 19, 29] (**Figure 1A,B**). We used a pupil index that removes the parts of pupil size explained by the visual stimulus and by slow drift (Supplementary Figure S1A). Frames were classified as high arousal if the pupil index was at or above the scan median, and as low arousal if it was below the median. We analyzed stretches of one state lasting at least 10 s. We matched the two states for visual stimulus and session time (Methods, Supplementary Figure S1B,C).

**Figure 1:**
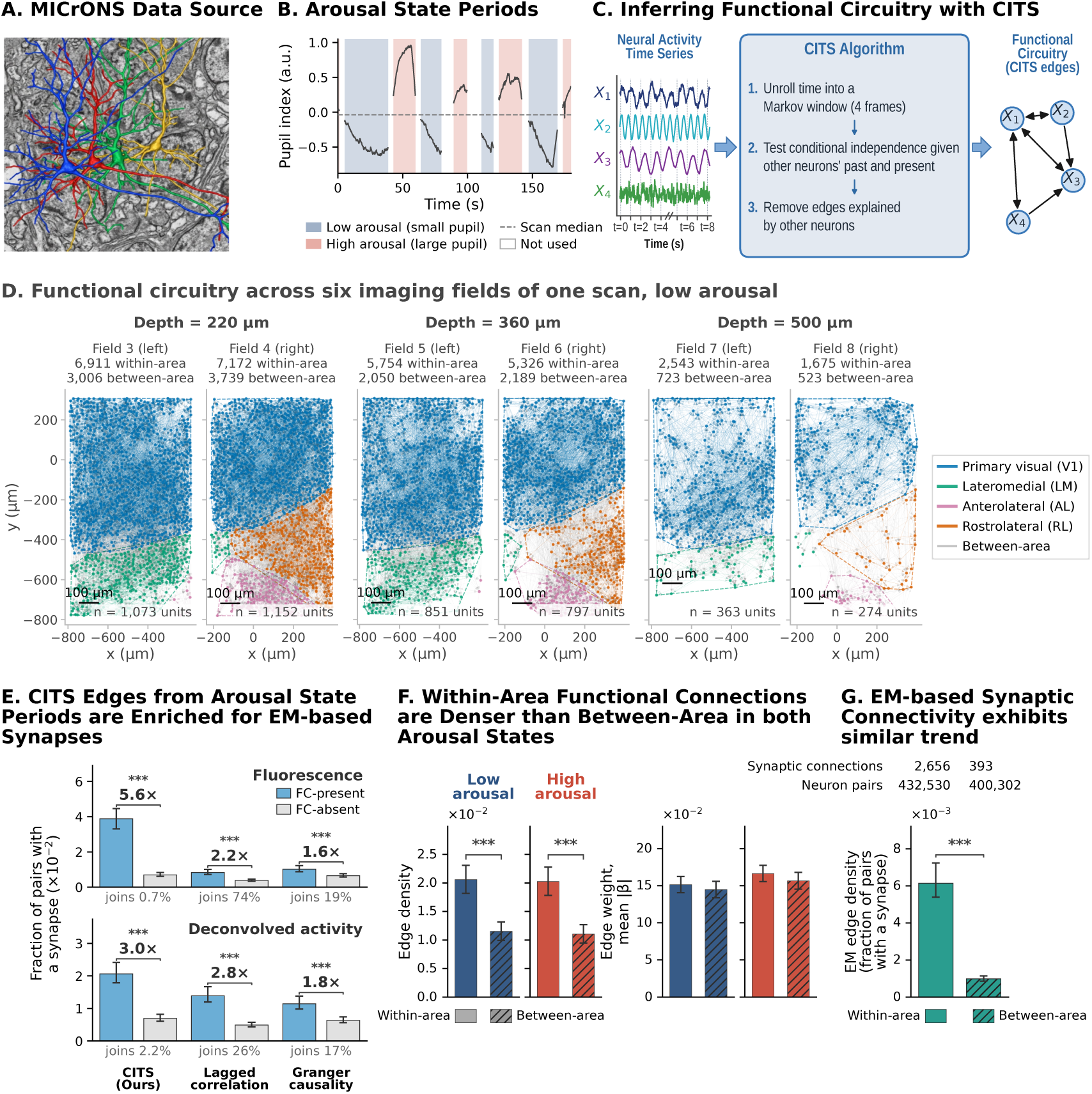
Data-driven inference of functional circuitry from large-scale measurements in mouse visual cortex. **(A)** The MICrONS dataset combines large-scale calcium imaging with dense electron-microscopy (EM) reconstruction [23]. **(B)** Arousal state periods defined by median split of the pupil index (computed per session and scan) [18, 19]. The example shows three minutes of session 7, scan 3. Shaded periods are stretches of one state lasting at least 10 s. **(C)** The CITS algorithm infers directed functional connections with statistical causal guarantees [9, 27]. **(D)** CITS-inferred microscale functional circuitry graphs for six imaging fields of one scan (session 4, scan 7) in low arousal. **(E)** Fraction of neuron pairs with an EM synapse, among pairs joined (FC-present) and not joined (FC-absent) by each method. Graphs of both arousal states are pooled, over *n* = 44 EM-co-registered fields. Top, fluorescence, with the same periods, blocks and settings as the deconvolved activity below. Bottom, deconvolved activity, used in all other panels. Brackets give the risk ratio of FC-present over FC-absent pairs. Under each sub-panel is the share of neuron pairs each method joined, because sparser edge sets tend to be more enriched. The panel assesses whether each map is consistent with synaptic anatomy, not the relative accuracy of the methods. On activity, 377 of 18,282 CITS-joined pairs had a synapse, and 240 of 6,191 on fluorescence. Error bars show 95% CIs, and stars mark *p <* 0.001 against 1. **(F)** Within-area functional connections are denser than between-area connections in both arousal states. Bars show directed edge density and mean *|β|*, with the mean and 95% CI over 12 scans. Asterisks mark *q <* 0.001, corrected for multiple comparisons. **(G)** Anatomical synaptic connectivity shows the same within *>* between organization as the functional circuit (*p <* 0.001). between 0.012 [0.010, 0.013], *q* = 0.00049). Both differences held in all 12 scans. Aligned to this, the EM synaptic connectivity exhibits the same phenomenon. Across areas, within-area neuron pairs are significantly more likely to be synaptically connected than between-area pairs in the synaptic connectome (0.61% versus 0.10% of ordered co-registered pairs, *p <* 0.001, Figure 1G). Both functional and synaptic connections were concentrated among nearby neurons. Comparisons of areas, layers and states used the 12 scans as samples and were corrected for multiple comparisons (Methods).

To infer directed functional circuitry from the calcium traces, we employed CITS [9, 27], a non-parametric statistical causal-inference method for neural time series (**Figure 1C**, also see **Methods**). It outperforms other approaches, such as Granger causality and PC-based methods, in reconstructing neural circuits [10, 27, 32], is scalable to the sizes needed for this dataset, and on this dataset its inferred connections are enriched for EM synapses (**Figure 1E**). CITS keeps an edge only when two neurons remain dependent after conditioning on sets of other neurons’ past and present activity. It yields same-time edges within one frame and lagged edges across one frame. We analyzed 44,987 active units in 96 imaging fields from 12 scans (Methods). Field-level CFC estimates were then compared between arousal states.

CITS was run on the data of each arousal state independently, yielding a directed functional connectivity graph over all neuron pairs within each imaging field per arousal state (**Figure 1D**). We summarized this graph at the area level by computing *edge density* for each directed area pair, defined as the fraction of all directed neuron pairs within that area pair for which CITS inferred a connection (**Figure 1D,F**). Each edge carries a weight, the standardized regression coefficient of the target neuron on its parents (Methods). Edge weight, also called edge strength, is the mean absolute coefficient, mean *|β|*, over the edges of a state’s graph.

We compared the functional graphs with the co-registered EM reconstruction in 44 imaging fields (**Figure 1E**). This comparison asks whether the inferred map is consistent with synaptic anatomy. It is not designed to benchmark inference methods (Discussion). At the single-connection level, a directed functional connection is roughly 3.0-fold more likely to have an EM synapse than a functionally unconnected pair (risk ratio 2.99, 95% CI [2.59, 3.31]). Edges of lagged correlation and conditional Granger causality on the same neurons were also enriched, 2.8-fold and 1.8-fold (risk ratios 2.84 and 1.81, **Figure 1E**). To test whether the enrichment depends on deconvolution, we repeated the comparison on the fluorescence traces of the same periods. The enrichment held there as well (**Figure 1E**, top).

Consistent with the local bias of cortical anatomy, intra-areal functional connections were denser than inter-areal connections in both arousal states (**Figure 1F**). In high arousal, within-area edge density (mean 0.020, 95% CI [0.018, 0.023]) substantially exceeds between-area density (0.011 [0.009, 0.013], *q* = 0.00049). The same pattern holds in low arousal (within 0.021 [0.018, 0.023],

### 2.2 Functional circuit organization varies across visual areas in both arousal states and is aligned to the synaptic connectome

The mouse visual cortex comprises multiple areas (V1, LM, AL, RL) with distinct functional specializations and hierarchical positions [24, 30]. We characterized the microscale functional circuitry organization across all 16 directed area pairs separately for each arousal state. This revealed a heterogeneous organization on top of an intra-areal dominant backbone (**Figure 2A,B**). In both arousal states, AL exhibits the highest within-area density, above LM, RL and V1 (*p* = 0.002). AL’s within-area mean edge density in low arousal is 0.062 (95% CI [0.045, 0.078]) and in high arousal is 0.062 [0.045, 0.079]. In contrast, V1 has the lowest with 0.020 [0.017, 0.022] in low arousal and 0.019 [0.017, 0.021] in high arousal, while LM and RL have intermediate edge densities (0.024–0.037). This is consistent with a higher-order area having stronger local recurrence [25,33,34]. Between-area connections are markedly sparser, with the AL*↔*RL axis forming the densest inter-area pathway in both states. RL*→*AL has a mean edge density of 0.026 [0.019, 0.033] in low arousal and 0.025 [0.017, 0.033] in high. AL*→*RL has a mean edge density of 0.026 [0.019, 0.033] in low arousal and 0.026 [0.017, 0.034] in high. These are substantially above all other between-area pairs (*p* = 0.002).

**Figure 2:**
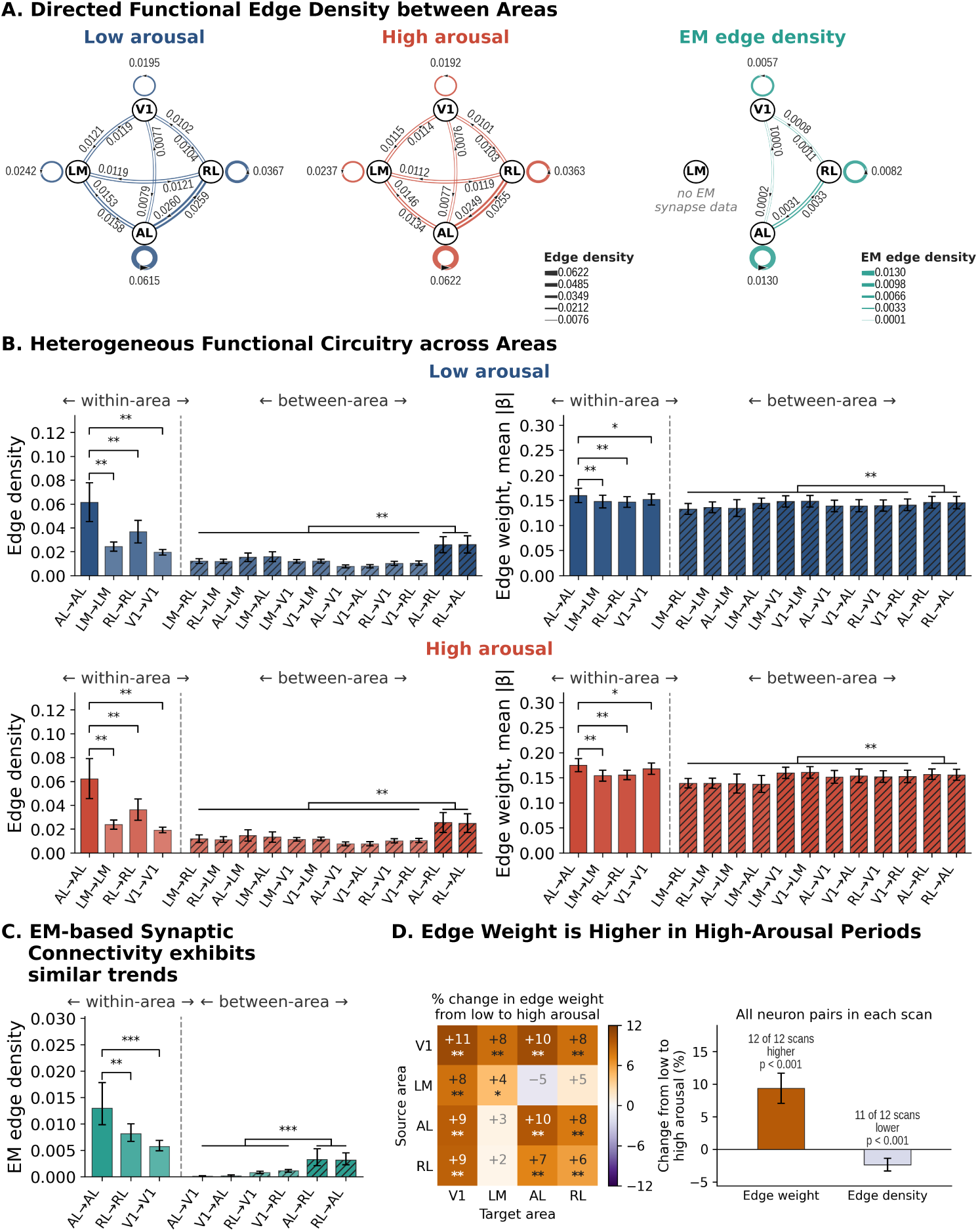
Areal functional circuitry is heterogeneous but consistent with EM synaptic connectivity, with edge weight higher in high arousal. (**A**) Graphical representation of the areal functional circuit edge density (low and high arousal) with EM synaptic connection density, showing that functional and anatomical circuits share the same intra-areal-dominant organization with an AL*↔*RL between-area backbone. LM had too few co-registered pairs for EM. **(B)** Edge density and edge weight (mean *|β|*) for all 16 directed area pairs for low arousal (top row) and high arousal (bottom row). Bars show the mean and 95% CI over 12 scans. Brackets mark significant differences between area pairs, corrected for multiple comparisons (* *p <* 0.05, ** *p <* 0.01). **(C)** Electron-microscopy (EM) synaptic connectivity provides the fraction of co-registered neuron pairs connected by an EM synapse per directed area pair, spanning *n* = 44 EM-co-registered imaging fields with reconstructed synapses. Area pairs involving LM are not shown, because each had at most 865 co-registered neuron pairs and at most one with a synapse. Error bars show 95% CIs. Brackets mark significant differences, corrected for multiple comparisons (** *p <* 0.01, *** *p <* 0.001). **(D)** Percentage change in edge weight from low-arousal to high-arousal periods for each ordered area pair. Stars mark significant changes, corrected for multiple comparisons (* *q <* 0.05, ** *q <* 0.01, *** *q <* 0.001), and faded squares are not significant. Bars on the right show the change in edge weight and edge density over all neuron pairs, pooled over the fields of each scan. Error bars show 95% CIs. Labels above the bars give the number of scans, out of 12, that changed in the direction of the bar.

AL had higher within-area edge weight than LM, RL and V1 in both states (*p ≤* 0.046, **Figure 2B**). Among individual between-area edges, the AL*↔*RL pathway has relatively strong between-area weights (*p ≤* 0.009 against the other between-area pairs). RL*→*AL has a mean edge weight of 0.145 [0.133, 0.158] in low arousal and 0.156 [0.145, 0.167] in high. These patterns reveal that while functional circuitry in both arousal states is intra-areal dominant, there is an inter-area backbone dominated by the AL*↔*RL pathway.

Between arousal states, this areal organization was largely preserved in terms of both areal edge density and strength. Edge strength was significantly higher in high arousal in 12 of the 16 area pairs (**Figure 2D**).

The same organization appears in the co-registered synaptic connectome (**Figure 2A,C**). Both the functional and synaptic circuits are intra-areal dominant with an AL*↔*RL between-area back-bone. Functional edge density and synaptic density were rank-correlated across area pairs, with Spearman *r* = 0.93 in low arousal and *r* = 0.95 in high arousal (*p <* 0.001) over the nine area pairs without LM. Pairs with higher functional density tended to have higher synaptic density, while the absolute densities differ in scale. LM was excluded for sparse co-registration.

### 2.3 Edge weight is higher in high-arousal periods

While the functional circuit organization was similar between arousal states, edge weight was higher in high-arousal than in low-arousal periods (**Figure 2D**). Over all neuron pairs, mean *|β|* was 9.4% higher (95% CI 7.1% to 11.7%, *q <* 0.001). It was higher in all 12 scans, by 2.7% to 14.2%, and in all five sessions (*p* = 0.00049, the smallest value possible with 12 scans). Resampling sessions, scans and fields gave a 95% CI of 6.2% to 11.5%. The increase was consistent within each scan, but smaller than the differences between scans. It was therefore significant only when the two states were compared within the same scan (without pairing, *p* = 0.059, Supplementary Table S1). The increase was 9.8% within areas and 8.2% between areas. The difference was robust to multiple controls (Supplementary Table S2). With the running controls added, the increase was 9.8%. Also excluding saccade windows gave 8.6% on 11 scans. Further removing movement and brightness signals from activity gave 8.4%. Every design showed the increase in every scan. The 9.4% increase in edge weight occurred although activity was about 10% lower in high-arousal periods. Among neurons with nearly unchanged rates, edge weight was also 9.1% higher on average in high-arousal periods, and higher in all 12 scans.

Twelve of the 16 ordered area pairs showed a significant increase, from 4.5% to 10.6% (*q <* 0.05). V1*→*V1 showed the largest increase. The four remaining pairs, all between LM and AL or RL, were small and uncertain, from *−*4.5% to +4.6%. The 12 significant area pairs were also robust to removing movement signals and imaging-brightness signals from each neuron’s activity.

Edge density showed no consistent change between states. It was 2.4% lower with the fields of each scan pooled (95% CI *−*3.3% to *−*1.4%, *q* = 0.0015, **Figure 2D**, right). Pooling gives larger fields more weight. The decrease was confined to the larger half of each scan’s fields (*−*2.9%, against +0.8% in the smaller half, difference *p* = 0.0015).

### 2.4 Cell-type and laminar functional circuitry resembles the synaptic connectome

Cortical circuitry is organized along multiple dimensions such as cell type, laminar position, and spatial extent [24, 35, 36]. Having characterized how microscale functional circuitry is organized in visual areas in arousal states, we next focused on its organization in cell types and layers.

We constructed aggregate cell-type functional circuitry networks for each arousal state, aggregated over *n* = 44 imaging fields with co-registered EM morphological cell type labels (see **Figure 3A**). Across the 84 drawn cell-type pairs, functional and EM densities were positively correlated (Spearman *r* = 0.28 for the low-arousal graph and *r* = 0.33 for the high-arousal graph). Cortical layers differ in their connectivity patterns and computational roles. Layer 4 receives thalamic input, layers 2/3 perform local and inter-areal integration, layer 5 provides cortical output, and layer 6 maintains corticothalamic feedback projections [37–39]. We characterized the layer organization of microscale functional circuitry in each arousal state (**Figure 3B**).

**Figure 3:**
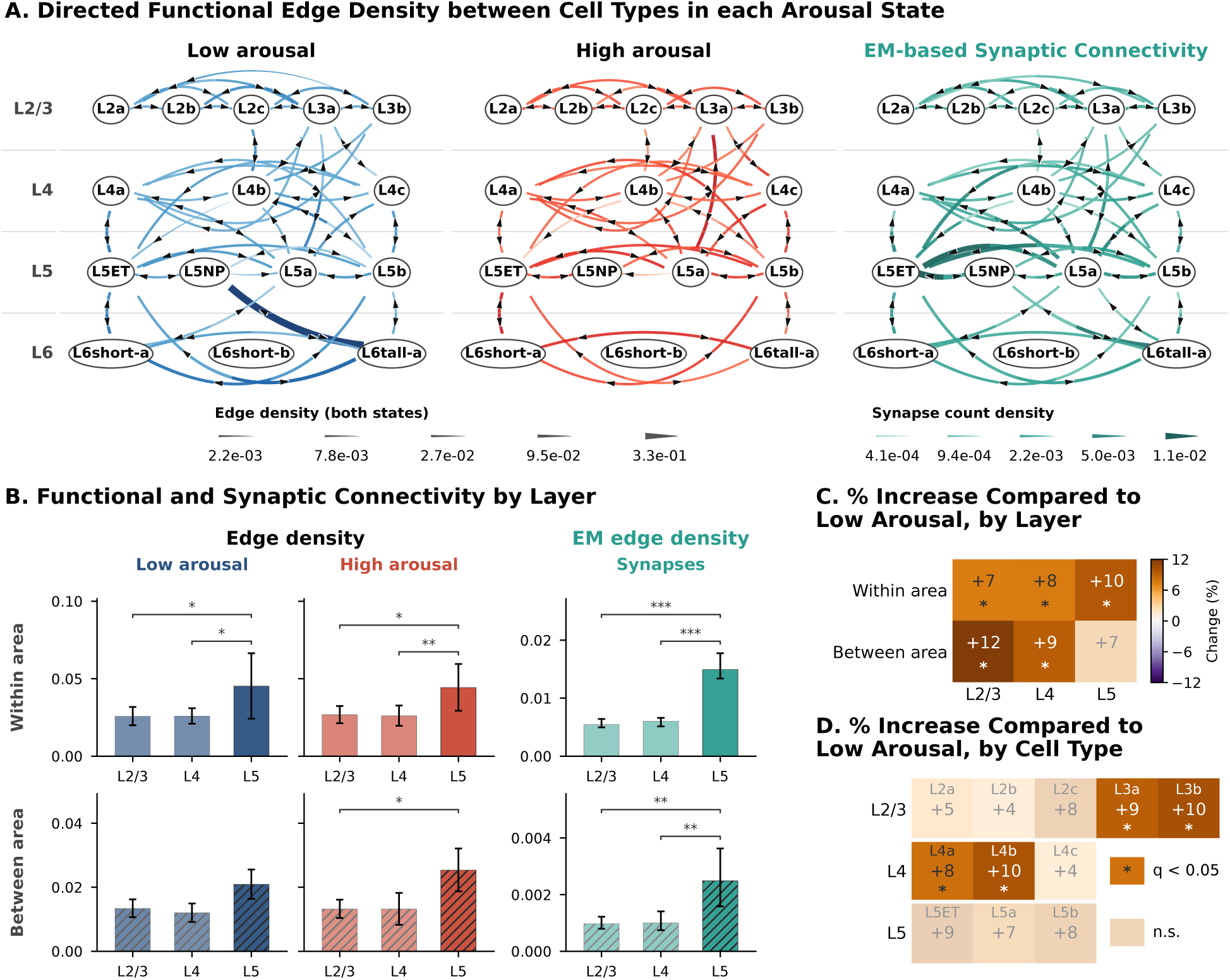
Cell-type and laminar functional circuitry is consistent with EM synaptic connectivity across arousal states, with edge weight higher in high arousal. (**A**) Cell-type functional circuitry networks (edge density) for low (left) and high (mid) arousal, shown together with corresponding EM synaptic connectivity (right). Excitatory cell types supported by at least three co-registered fields are included. Edge density comes from the *n* = 44 fields with co-registered EM morphological cell-type labels. In the functional graphs, layer-6 nodes rest on only 40 co-registered neurons in 10 of the 44 fields. (B) Within-area and between-area edge density per arousal state at layers 2/3, 4, and 5, with EM synaptic connectivity across layers alongside. Layer 6 is omitted here due to too few layer-6 neurons co-registered per field for a reliable field-level estimate. Brackets mark significant differences between layers, corrected for multiple comparisons (* *p <* 0.05, ** *p <* 0.01, *** *p <* 0.001). **(C)** Change in functional connection strength (high *−* low arousal) for each layer group, in percent of the low-arousal value. Stars mark *q <* 0.05, corrected for multiple comparisons. Faded squares are not significant. Each value is the mean over seven to 11 scans, depending on the layer group. **(D)** The same change for the connections of each excitatory cell type, incoming and outgoing together. Stars mark *q <* 0.05, corrected for multiple comparisons. Faded squares are not significant. Each value is the mean over seven to 11 scans, depending on the cell type. equal field weights and in both halves of the data (Methods). We treat the cell-type results as exploratory.

Layer 5 was the densest layer in the EM connectome. Within area, connection probability was 0.0150 with 95% CI [0.0134, 0.0177] in layer 5, versus 0.0055 [0.0050, 0.0064] in layer 2/3 and 0.0061 [0.0051, 0.0066] in layer 4. Between area, it was 0.00249 [0.00158, 0.00363] in layer 5, versus 0.00098 [0.00079, 0.00122] in layer 2/3 and 0.00101 [0.00075, 0.00141] in layer 4.

The microscale functional circuit showed the same within-area layer-5 prominence. Within each arousal state, functional circuitry was densest in layer 5 with a mean edge density of 0.045, 95% CI [0.024, 0.066] in low arousal and 0.044 [0.029, 0.059] in high (*p ≤* 0.023 against layers 2/3 and 4). Layer 2/3 (low 0.026 [0.020, 0.032], high 0.027 [0.021, 0.032]) and layer 4 (low 0.026 [0.021, 0.031], high 0.026 [0.020, 0.033]) were sparser and similar. Between areas, layer 5 was denser than layer 2/3 only in high arousal (*p* = 0.031). Between arousal states, within-area and between-area edge strength were higher in high than in low arousal in all six layer groups, by 7.2% to 11.9% (**Figure 3C**). This increase was significant in five groups (*q ≤* 0.026), but not for layer 5 between areas (*q* = 0.11).

While the overall organization is shared between arousal states, the edge weight of each excitatory cell type’s connections was higher in high-arousal periods for all 11 types shown, by 4% to 10% (**Figure 3D**). Four of the 12 tested types, L3a, L3b, L4a and L4b, had *q <* 0.05 (Supplementary Table S1). All four held across sessions, after removal of movement and brightness signals, with

### 2.5 Stimulus-driven predictive correlation covaries with the strength of microscale functional circuitry

A key question in systems neuroscience is whether circuit-level organization is associated with a measurable difference in the brain’s ability to represent sensory stimuli [40]. To address this, we computed predictive correlation, i.e., how well a stimulus-driven model predicts each neuron’s response, as a proxy for sensory encoding capacity [41]. To obtain predictive correlation, we built a predictive model using the winning architecture from the Sensorium 2023 NeurIPS competition [42]. We trained the model on MICrONS data with seven-fold cross-validation and evaluated on held-out folds (**Figure 4A**, see **Methods**). Using the predictive model, we computed predictive correlation for each neuron, as the correlation between the model-predicted and the actual neural responses, for each arousal state. The analysis covers 111,378 neuronal units across 12 scans.

**Figure 4:**
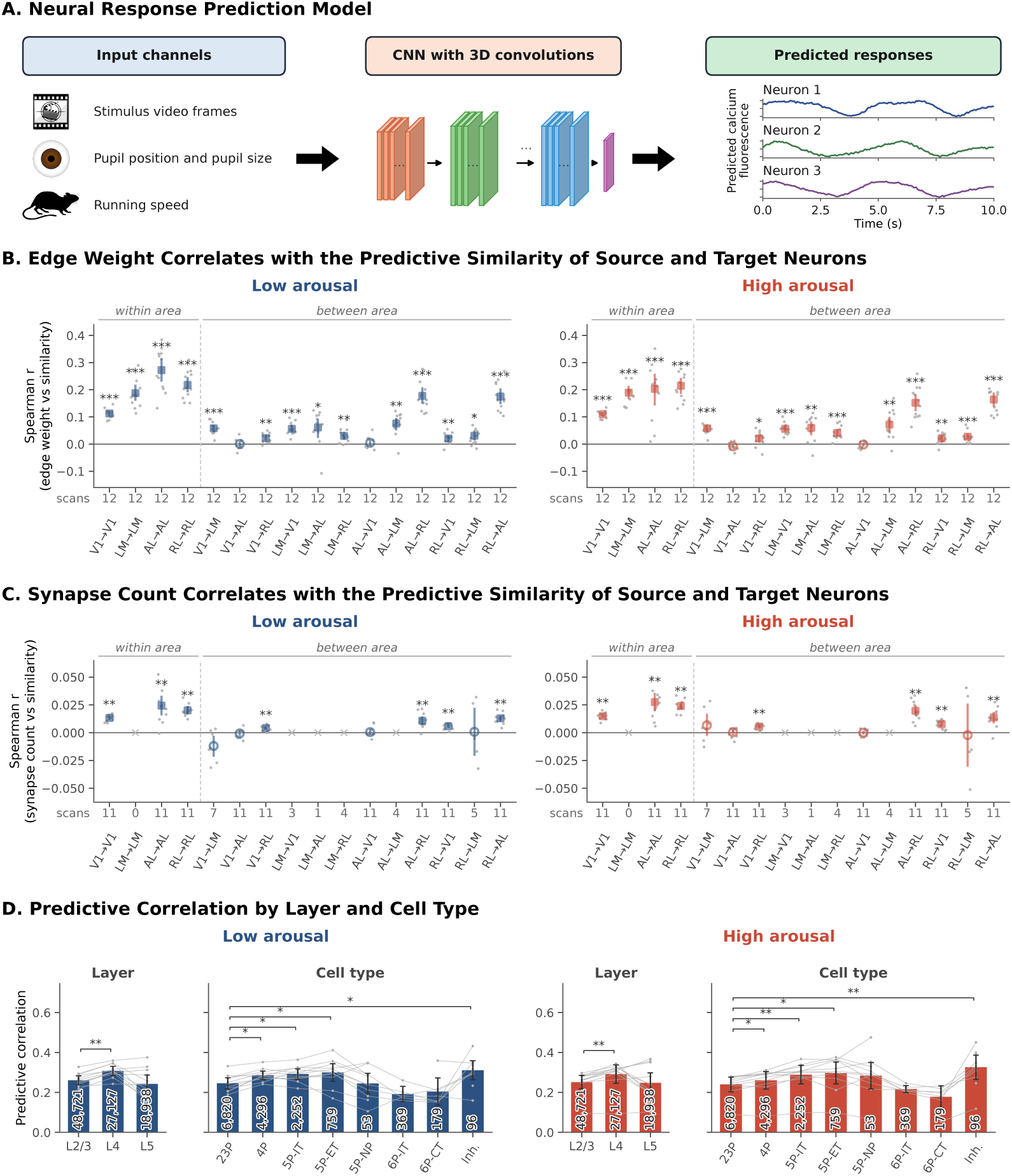
Neural sensory encoding performance aligns with functional circuitry and EM synaptic connectivity in both arousal states. (**A**) Schematic of the neural response predictive model used to compute predictive correlation, from the visual stimulus, pupil position, pupil size and running speed [42]. **(B)** Functional circuitry weight correlates with encoding similarity, as indicated by Spearman correlation between CITS-inferred microscale functional circuitry strength and predictive correlation profile similarity, per directed area pair and arousal state (mean *±* 95% CI over 12 scans). **(C)** Synaptic connectivity count correlates with encoding similarity, as indicated by Spearman correlation between EM synapse count and predictive correlation profile similarity, per directed area pair and arousal state (11 scans). Crosses mark pairs with fewer than five scans. We did not test these pairs or plot their scan values. **(D)** Predictive correlation by cortical layer and cell type across arousal states. The panel uses all imaged neurons of the 12 scans, and numbers in the bars give neurons. Bars show the mean and 95% CI over scans, and gray lines show single scans. Asterisks indicate significant differences, corrected for multiple comparisons. For cell types, we show significance for seven pairs only. These are 23P against 4P, 5P-IT, 5P-ET and inhibitory cells, 5P-IT against 5P-ET and 6P-IT, and 6P-CT against inhibitory cells.

CITS-inferred functional circuitry strength was positively correlated with predictive correlation profile similarity (predictive similarity) in both arousal states (**Figure 4B**). Here profile similarity measures how similarly two neurons’ encoding performance varies across trials. In other words, neurons with stronger functional connections exhibit more correlated encoding performance across trials. The mean Spearman correlation was 0.183 within areas and 0.063 between areas for the low-arousal graph. It was 0.165 and 0.056 for the high-arousal graph. In each state, the correlation was significantly positive in 14 of 16 ordered area pairs (*q <* 0.05, Methods), all except V1*→*AL and AL*→*V1. In both arousal states, the relation of functional circuitry strength to encoding similarity was significantly stronger for within-area pairs, particularly AL*→*AL and RL*→*RL, than between-area pairs (*p* = 0.00049). This held in all 12 scans. The relation did not differ significantly between the two graphs (within areas, difference *−*0.017, 95% CI [*−*0.035, 0.003]).

The EM synaptic connectivity count showed the same positive relationship with encoding-profile similarity (**Figure 4C**), resolvable in fewer pairs given the sparser co-registered synaptic sampling. Furthermore, across area pairs, the functional and synaptic relationships to encoding similarity followed the same pattern (**Figure 4B,C**). The same area pairs ranked highest in both, with the within-area pairs AL*→*AL and RL*→*RL at the top. The synaptic correlation was significantly positive in seven area pairs in both states, including the three testable within-area pairs, with means of 0.004 to 0.027.

Predictive correlation varied by cortical layer and cell type in both arousal states (**Figure 4D**). Among layers, L4 showed the highest mean predictive correlation in both low and high arousal. L4 has mean predictive correlation of 0.308, 95% CI [0.285, 0.331] in low arousal and that of 0.292 [0.245, 0.338] in high arousal, significantly exceeding L2/3 in both states (*q <* 0.01). Among excitatory cell types, layer-5 extratelencephalic neurons (5P-ET) showed the highest predictive correlation in both states with mean predictive correlation of 0.300 in low arousal and that of 0.297 in high arousal. Layer-2/3 pyramidal cells (23P) had lower predictive correlation than 4P, 5P-IT and 5P-ET cells in both states. Layer-6 intratelencephalic and corticothalamic subtypes (6P-IT, 6P-CT) showed the lowest predictive correlations (6P-IT low 0.192, 6P-CT low 0.204).

## 3 Discussion

This study characterizes microscale functional circuitry in mouse visual cortex at single-neuron resolution across arousal states. Functional circuitry is mapped at different scales of organization spanning area-level topology, cell-type and laminar motifs, and sensory encoding. At each scale of organization, we compared the resulting functional circuitry with the co-registered synaptic connectome.

The CITS-inferred directed functional connections align in areal, laminar, and cell-type organization with the co-registered synaptic connectome. This structure-function correspondence reflects a different facet than what was previously reported. Earlier studies in mouse visual cortex related synaptic connectivity to response similarity [13, 14]. They found that neuron pairs with correlated visual responses and similar tuning are connected at higher rates and through stronger synapses. However, such similarity is undirected and defined by the stimulus. In contrast, we compared directed microscale functional circuitry inferred from temporal dependencies in neural activity to the synaptic connectome.

The EM comparison assesses whether an inferred map is consistent with synaptic anatomy, rather than benchmarking inference methods. Edges of all three methods were enriched for synapses on both traces (**Figure 1E**). Enrichment is not suited to ranking methods, because it depends on how many pairs each method joins. Synapse counts also cannot isolate the indirect links that CITS is designed to remove. Our earlier study benchmarked CITS against known connectivity in simulated networks [27]. There, CITS outperformed pairwise and conditional Granger causality and PC-based methods. The network-wide edge-weight increase in high arousal appeared with all three methods (Supplementary Table S2).

The finding that within-area connections dominate functional circuitry in both arousal states is consistent with the cortical column’s information processing [24, 43]. Dense recurrent connectivity within areas supports local computation. Sparse between-area connections carry the output of this local processing across the visual hierarchy [30, 44]. AL exhibits the highest within-area connection density in both states. This may reflect its role as a higher-order visual area with stronger local recurrence [25, 33, 34]. The dominance of the AL*↔*RL inter-area axis is consistent with the lateral, non-hierarchical connectivity between AL and RL identified in experimental studies [45]. Unlike feedforward V1*→*LM projections, the AL*↔*RL connection links areas at similar hierarchical levels [24]. This is consistent with lateral integration within the dorsal motion-processing stream rather than hierarchical feedforward transmission [33, 44].

Because few neurons of each cell type are co-registered per field, individual subtype edges are small-sample and are not interpreted individually. The layer-level analysis has more neurons per field than any single cell type, allowing significance testing. Within-area functional density was highest in layer 5, significantly above layer 4 in both states. This matches the EM synaptic connectome, where layer 5 is the densest layer. Paired recordings in rat visual cortex also found dense, often reciprocal connections among layer-5 pyramidal neurons [46].

Pupil-linked arousal varies on fast and slow time scales. Rapid pupil dilations accompany phasic noradrenergic activity, and longer-lasting dilations accompany sustained cholinergic activity [28]. Earlier work found less correlated population activity during rapid dilations [19] and lower noise correlations during heightened arousal [47]. Our analysis addresses the slower component. We compared sustained states of at least 10 s and measured conditional dependence between neurons, including shared stimulus responses. In these sustained states, CITS edge weight and pairwise correlation were both higher in high-arousal periods (**Figure 2D**, Supplementary Table S2). Fast and sustained components of arousal may relate to cortical coordination in different ways.

The increase was positive in every layer group and in every excitatory cell type shown (**Figure 3C,D**), so it was not confined to one laminar circuit. The organization was preserved, and edge density showed no consistent change. This pattern is consistent with a gain change on existing interactions, not the recruitment of new ones. The data do not identify the source of the difference. Candidate sources are neuromodulatory state [28], movement-related input [48, 49] and visual drive that differs with pupil size [50]. Firing rate is an unlikely source, because the difference persisted in neurons with stable rates. Running is also an unlikely source, because excluding running periods slightly increased the difference. Eye movement remains a candidate. Within each state, edge weight was on average 7.4% higher during faster eye movement. Eye speed was also higher in high-arousal periods. Our data cannot separate eye movement from arousal. A difference in imaging noise between states remains possible, because lower noise would raise estimated edge weights. The 2.9% density decrease in the larger fields may be the more accurate estimate. Larger fields record more of the neurons that could explain a dependence, so CITS can remove more indirect edges.

The positive correlation between CITS functional circuitry weight and predictive correlation profile similarity (**Figure 4B**) shows that stronger functional connections link neurons with more similar sensory encoding performance. This parallels the like-to-like synaptic wiring of mouse visual cortex described above [13, 14]. This relationship holds across both arousal states and is significant for all within-area directed pairs. The synaptic measure points the same way. EM synapse count also correlates positively with encoding-profile similarity (**Figure 4C**), agreeing in sign with the functional measure in the seven area pairs where it was significant. Because co-registered synaptic pairs are sparse, this relationship is resolvable in only a subset of area pairs anatomically, whereas the functional measure recovers it in 14 of 16 area pairs. The functional circuit thus provides a denser readout of a relationship that the anatomy samples only partially.

The variation in predictive correlation across layers and cell types (**Figure 4D**) is consistent with known hierarchical specialization of visual cortex [24, 30]. L4 carries the highest predictive correlation among all layers in both states. L4 is the primary recipient of thalamocortical input and is therefore most tightly coupled to the sensory stimulus. Among excitatory cell types, 5P-ET neurons show the highest predictive correlation. Layer-5 extratelencephalic neurons are a principal cortical output class. Their high stimulus predictability is consistent with reliably stimulus-driven activity, matching their role in relaying visual information to subcortical targets [24]. At the other extreme, L6 excitatory subtypes (6P-IT, 6P-CT) show the lowest predictive correlation.

This study has several methodological boundaries that motivate future work. First, pupil size is a well-established proxy for arousal [18, 19, 29], but does not capture the full complexity of arousal-related neuromodulation. Pupil size also correlates with locomotion, and the two co-vary in this mouse. Running was sparse overall, but the running episodes that occurred tended to coincide with high-arousal, large-pupil trials. The edge-weight increase held after we controlled for running, excluded saccade windows and regressed movement signals from neural activity (Supplementary Table S2). Eye speed was higher in high-arousal periods. Orofacial movements were not measured. Disentangling pupil size from eye movement would require recordings with greater within-state eye-movement variation. Future studies combining optogenetic manipulation of neuromodulatory systems with functional imaging would provide more direct causal evidence for the mechanisms proposed here. Relevant targets include locus coeruleus and basal forebrain. Second, our analysis focused on visual cortex. Extending these findings to other sensory and association areas would clarify whether the edge-weight difference between states is a general property of neocortex or specific to sensory hierarchies [51, 52]. Third, causal sufficiency is a standard assumption in statistical causal inference that is challenging to satisfy in vivo. All common causes of recorded neurons must be observed [53]. Unobserved inputs such as thalamic drive and neuromodulatory fluctuations cannot be fully excluded. CITS partially addresses this by testing, for each connection, whether it holds after controlling for subsets of other neurons’ activity in the lag window. When a recorded neuron is driven by the same unobserved input as the pair being tested, controlling for it absorbs part of that input’s influence. This can weaken or remove the spurious edge [53]. Because functional and synaptic connections were both concentrated among nearby neurons, part of the synapse enrichment of functional edges may reflect proximity alone. Nearby neurons can be functionally connected without a synapse, through shared unrecorded input or mixing of their calcium signals. Perturbation experiments targeting specific cell types would provide direct validation of the inferred connections. At 6.3 Hz, the direction of same-time edges cannot be identified. We interpret only aggregate measures, not single edges. Fourth, the cell-type classification used here aggregates many subtypes. Future work with more granular cell-type identification (e.g., specific interneuron subtypes) could reveal more detailed mechanisms [35, 36, 54]. Directed inter-laminar (cross-layer) connectivity is a separate question, additionally constrained by the single-plane imaging design. Only 18 of 43 fields captured neurons from two layers in the same area, four to 11 per layer pair. Cross-layer flow therefore could not be reliably assessed at the field level. Each field is one plane, so layer comparisons are also field comparisons, and edge density depends on field size, as in any network [55]. Finally, this analysis was limited to one awake, head-fixed mouse viewing natural movies and synthetic parametric videos. A held-out session of the same animal failed its pre-specified quality gates, so no replication is available. The analyses reported here were specified after an inconclusive pre-specified analysis (Methods). How these findings extend to other behavioral states (e.g., sleep, active locomotion) remains an open question [49, 56]. Despite these limitations, the directed circuitry inferred here yields specific, testable predictions to guide targeted perturbation experiments.

## 4 Methods

### 4.1 Data Source

This study is observational. We did not manipulate arousal or any other state. We used the MICrONS dataset (Minnie65) [23], which pairs a dense electron-microscopy reconstruction with two-photon calcium imaging (GCaMP6s [57]) of the mouse visual cortex. The dataset comprises detrended calcium fluorescence traces from 146,388 units across 16 session-scans, of which 15,434 were co-registered with the EM volume. We retained the 13 session-scans with intact pupil recordings, excluding 3 with corrupted pupil data. Twelve met the design criteria and passed the label-validity gate (below). We analyzed these 12 scans from five sessions, with 96 imaging fields. For the predictive correlation analysis, the unit of observation is a session-scan-unit combination. There were 111,378 units across these 12 scans, a mean 9,282 units per session-scan (range 8,395–12,675). Stratified analyses (e.g., by brain area or cell type) use annotated subsets of these units and report the corresponding sample size in each case.

### 4.2 Arousal State Classification

Arousal denotes the level of alertness of an awake animal, from drowsiness to heightened vigilance [58]. Noradrenaline from the locus coeruleus and acetylcholine from the basal forebrain can modulate cortical state across these levels [58]. In mice, sensory detection is best at intermediate arousal [18]. Pupil fluctuations track rapid changes of cortical state during quiet wakefulness [19]. They also follow noradrenergic and cholinergic activity in cortex [28] and the firing of locus coeruleus neurons in monkeys [29]. Arousal often co-occurs with locomotion, but the two have distinct correlates in cortical activity [47].

We defined arousal states based on pupil size, a well-established proxy for arousal [18,19,29,59]. We measured pupil size as the natural logarithm of the major radius of the fitted pupil ellipse. We removed the part of this signal explained by the visual stimulus. We fitted log major radius by least squares on all frames with a valid pupil fit. The predictors were stimulus type and the luminance, contrast and motion energy of the movie at lags of 0 to 20 frames. We did not remove running or other movement from the index (see Sensitivity to the controls). The pupil index is the residual minus its centered 10-minute moving median (**Figure 1B**, Supplementary Figure S1A). Pupil changes lag the neuromodulatory activity they track [29]. We therefore labeled each neural frame with the pupil index six frames (0.95 s) later. A frame is high-arousal if its index is at or above the scan median, and low-arousal otherwise. Splitting at this scan median accounts for scan-to-scan variation in pupil size. The predictive-correlation analysis (below) uses whole trials, so each trial needed one state label. We assigned it with a consistency criterion. A trial needed at least half of its frames in analyzed periods. We classified a trial as high-arousal only if *≥* 80% of its analyzed frames were high-arousal. We classified a trial as low-arousal only if *≥* 80% of its analyzed frames were low-arousal. We excluded trials that did not meet this consistency criterion from the predictive-correlation analysis. Scans had 65 to 129 low-arousal and 72 to 122 high-arousal trials. This approach ensures that each trial represents a relatively stable arousal state.

#### Exclusions, periods and blocks

We applied the same rules to both states. Frames without a pupil fit had no state label, so these gaps, padded by 1 s, ended a stretch of one state. Frames without a visual stimulus also ended a stretch, because the matching below uses stimulus covariates. So did the first and last 5 min of each scan, where the centered 10-minute moving median is incomplete. Brain motion corrupts the fluorescence signal. We therefore removed each frame whose field-mean frame-to-frame shift exceeded the median by five median absolute deviations, with its two neighbors. We did not exclude running or saccades. Saccades were detected as persistent steps in pupil-center position. A pupil sample taken during a saccade did not set a state label, because the pupil fit is unreliable then. We kept stretches of at least 10 s, so that each state was sustained. We trimmed 2 s from both ends of each stretch to avoid state transitions. We cut stretches into pieces at changes of stimulus type, so that each piece holds one stimulus type. We split each scan into two folds for held-out checks. Consecutive 60-s super-blocks alternate between the folds, and pieces also end at their borders. We dropped the first 6 s of each super-block, so frames of different folds are at least 6 s apart. This gap limits autocorrelated activity shared between the folds. Each piece is a grid of blocks of four consecutive frames (about 0.63 s). A block is one sample for CITS and one unit of matching. For each neuron, we subtracted the 10-minute moving median, which removes slow drift, and the mean within each stimulus type. We then centered activity within each piece.

#### Matching and gates

High-arousal and low-arousal periods could differ in what the mouse saw and in when they occurred in the session. Either difference alone could affect the estimated circuitry, so we matched the two states on both. Selection used behavioral, stimulus and imaging-quality signals and read no neural traces. Within each fold, we grouped blocks by stimulus type, by session-time quintile and by quintile of a screen-content score. This propensity score is the probability of high arousal predicted from the screen content by ridge-penalized logistic regression [60]. Its inputs were the block means of movie luminance, contrast, motion energy, the change in luminance over seven frames and position in the stretch. They also included 20 movie-content components, the first 20 principal components of image features from a pretrained ResNet-18 network [61, 62]. In each group, we kept equal numbers of blocks per state and thinned the larger state evenly in time. Matching within each fold keeps both folds balanced on their own. Each state kept 14,546 blocks (154 min) over the 12 scans, and 847 to 1,501 blocks per field (median 1,196.5). We did not match eye movement or running, because matching on them as well did not balance the states. Eye speed was higher in high-arousal periods in all 12 scans. Running occurred in 2.5% of high-arousal blocks and 0.2% of low-arousal blocks. A saccade window, from one frame before to 2 s after a saccade onset, overlapped 16.8% of high-arousal blocks and 5.3% of low-arousal blocks. The stricter designs below address these differences.

Two checks had to pass before we compared the states. The balance gate tested whether matching worked. Pooled over scans with block weights, every screen covariate had to differ between states by less than 0.1 standard deviations [63]. After matching, per-scan differences in luminance, contrast, motion energy and time in session were at most 0.09 standard deviations. Each stimulus type made up the same share of blocks in both states (Supplementary Figure S1C). The label-validity gate tested whether the pupil index has a neural correlate in each scan. A decoder trained on population activity in one fold predicted the block-mean index in the other fold. A scan passed if its decoding *R*^2^ exceeded that of all 20 pseudo-sessions, which pair its activity with the pupil index of another session. All 12 scans passed (*R*^2^ 0.31 to 0.59). This gate shows a neural correlate of the index, which movement-related activity could also supply.

### 4.3 Functional Circuitry Analysis

We estimated functional circuitry using CITS [9, 27], a nonparametric, time-aware causal-inference framework for neural time series. CITS infers connections from conditional-independence relations among neural activity, at both time-lagged and contemporaneous (zero-lag) intervals. Lagged connections are directed by time. Each edge’s weight is estimated by linear regression of each neuron on its inferred causal parents (see Algorithm details).

We applied a highly scalable instantiation of CITS to the deconvolved activity, testing conditional independence with the Fisher-z test. This instantiation is sensitive to monotone dependence. The autocorrelation the calcium indicator adds to each individual trace does not confound the inferred edges. The conditioning sets include the target neuron’s own past, so the indicator’s decay is absorbed by the autoregressive self-term. Consistent with this, the inferred edges are enriched for synaptic contacts (**Figure 1E**), which indicator dynamics alone would not produce. Activity is the deconvolved trace released with the dataset, computed by constrained deconvolution [64] in CaImAn [65].

Standard correlation cannot distinguish a direct connection between two neurons from an indirect one mediated by a third neuron. Pairwise Granger causality [66] addresses directionality but does not account for network-wide dependencies. Multivariate extensions do, but assume linear interactions between neurons [15]. The CITS framework makes no linearity or distributional assumption. It models the multivariate neural dynamics with a structural causal model of arbitrary Markov order [27]. It infers a directed edge from neuron *i* to neuron *j* when their activities remain statistically dependent at a time lag after conditioning on the past activity of the other neurons and of neuron *j* itself. This dependence can be assessed with nonparametric conditional-independence tests that capture nonlinear and non-Gaussian interactions. This removes indirect dependencies and yields a directed connectivity structure per field and arousal state.

#### Algorithm details

For neurons *i* and *j* with activity *X_i_*(*t*) and *X_j_*(*t*), CITS tests whether *X_j_*(*t*) remains dependent on *X_i_*(*t − τ* ) after conditioning on sets of other variables in a time-unrolled window, where *τ* is the time lag. An edge *i → j* is retained when this lagged dependence survives conditioning. With Markov order *τ* = 1, each sample is one block of four consecutive frames, and each neuron at each frame is one variable. CITS searches for the skeleton of conditional dependence among these variables with the PC-stable algorithm [53,67]. We did not cap the size of conditioning sets. We used a graphics-processing-unit implementation of the PC-stable search [68], extended to all conditioning-set sizes. On every graph we compared, it reproduced the central-processing-unit search exactly. We read out same-time edges between neurons at the last frame and lagged edges from the third frame to the last. The earlier frames serve only as conditioning context. We did not orient same-time edges. We estimated one graph per field and arousal state from all matched blocks of that state, with no split into folds. We combined these contemporaneous edges with the time-directed lagged edges to define the functional circuit. We then assigned edge weights following CITS [27]. Edge weights are standardized regression coefficients, computed from the correlation matrix of the window variables over the blocks of the state. For a lagged edge from *i* to *j*, we regressed *j* at the last frame on its lagged parents. The weight is the coefficient of *i* at the third frame. These include the past of *j* itself when the graph links it. For a same-time edge between *i* and *j*, we regressed *j* on *i* and on the lagged parents of *j*. The coefficient of *i* is the weight of *i → j*. The weight of *j → i* is defined in the same way. Each regression used at most 25 regressors. When a pair had both a same-time and a lagged edge, we used the same-time weights. No same-time edge had coefficients of opposite sign in its two regressions. We estimated weights on the same blocks that define the edges, so selection inflates absolute weights in both states alike. Positive weights indicate excitatory-like and negative weights inhibitory-like influence. We used a maximum lag of one frame (approximately 159 ms at 6.3 Hz [23]) and a significance level of *α* = 0.05 for the conditional independence tests. A one-frame lag matches the temporal resolution of calcium imaging, which is limited by indicator kinetics rather than by neural spiking.

#### Statistical causal inference guarantees

CITS recovers the correct connections consistently as the number of observed time points grows, under mild assumptions [10, 27]. The lagged edges are directed by time, and contemporaneous edges are identified up to Markov equivalence. These guarantees are nonparametric, i.e., without assuming any specific parametric form for neural dynamics. The partial-correlation instantiation used here is consistent for monotone dependence when the conditioning variables act linearly. Recovery rests on two standard conditions for causal discovery [53], which are 1) the directed Markov property, under which the graph structure matches the conditional-independence patterns in the data, and 2) faithfulness, under which every direct connection produces a dependency that survives conditioning, with no exact cancellations. Because the model includes temporally lagged variables, it captures inter-temporal relationships that contemporaneous-only methods miss. Faithfulness violations require exact cancellations, are generically measure-zero, and are unlikely to systematically bias the aggregate network patterns reported here. A third assumption, causal sufficiency, requires that no unobserved variable drives two recorded neurons. It is only partially met, because two-photon imaging samples a subset of the neurons in each field and none of the unrecorded cortical, thalamic, or neuromodulatory in-puts. Several features limit the resulting exposure. First, CITS conditions each candidate edge on the other recorded neurons and on each neuron’s own past. This absorbs common inputs from recorded cells and part of the influence of shared drivers reflected in their recent activity [27]. Second, for the lagged edges, CITS is provably robust to latent confounding under a first-order Markov structure without concurrent effects, recovering the correct lagged graph among observed variables [27]. The contemporaneous edges we additionally infer fall outside this guarantee and are the most exposed to unobserved common drive. Third, the primary comparisons are relative and within-field, contrasting arousal states or edge classes within the same neurons, so drivers shared across conditions largely cancel. Fourth, the inferred edges are enriched for synaptic contacts in the electron-microscopy reconstruction (**Figure 1E**), indicating that they carry information about anatomical connectivity. Residual confounding from unobserved sources remains possible, most plausibly for contemporaneous edges, so individual edges are interpreted cautiously and conclusions are drawn at the level of aggregate network organization. Performance on simulated neural circuits is described in Benchmarking below. The approach removes the indirect dependencies that confound correlation and assigns directionality to lagged edges. We analyze this statistically causal functional circuitry throughout.

#### Benchmarking

Prior work has benchmarked CITS on simulated neural time series against pair-wise (GC1) and conditional (GC2) Granger causality, kernel nonlinear Granger causality, the PC algorithm, the Time-Aware PC algorithm (TPC), and the state-of-the-art constraint-based time-series methods PCMCI^+^ and LPCMCI [10, 27, 32]. Recovery of the known ground-truth graph was scored by the combined score (CS = TPR *−* FPR). Across linear-Gaussian and nonlinear non-Gaussian autoregressive paradigms, CITS attained a mean CS of 0.999, exceeding the strongest baseline (TPC, 0.828) and the modern methods PCMCI^+^ (0.755) and LPCMCI (0.769). On biologically realistic GLM-coupled spiking networks with refractory self-history, CITS recovered within-neuron recurrence (mean CS 0.987), whereas the Granger family cannot represent self-connections and collapsed (CS 0.29–0.34). Through GPU parallelization, CITS scaled to networks of 1,000 neurons in about 33 seconds, a regime in which PCMCI^+^ and LPCMCI become computationally intractable. These results support CITS’s suitability for inferring directed functional circuitry at the scale of the recorded fields.

### 4.4 Data Processing Pipeline

#### Unit selection

We applied CITS separately within each of 96 imaging fields across 12 scans. For active-unit selection, we retained a unit if its deconvolved spike trace exceeded a value of one in more than 12% of frames during at least one stimulus type (Clip, Monet, or Trippy). CITS then estimated functional circuitry from the deconvolved activity of the retained units. Different stimulus types selectively engage different neurons. Taking the union across stimulus types captures every neuron with detectable stimulus-evoked activity in any part of the recording. Units below this threshold in all three stimulus types were excluded. Of the 111,378 units of the 12 scans, this yielded 44,987 active units (mean 469 per field) used for causal functional circuitry (CFC) inference. We counted a neuron imaged in more than one scan once in each scan.

#### CFC Matrix Computation

For each field, session, and scan, we applied CITS to deconvolved activity (as provided in the MICrONS processed dataset) separately for the matched blocks of each arousal state. This generated causal functional circuitry matrices (CFC matrices) where each entry (*i, j*) is the signed edge weight from neuron *i* to neuron *j*, the regression coefficient assigned by CITS (Algorithm details), or zero if no edge was inferred. Positive values indicate excitatory-like and negative values inhibitory-like influence. A same-time edge carries a separate weight in each direction (Algorithm details).

### 4.5 EM Synaptic Connectivity and Structure-Function Comparison

We linked functional units to EM-reconstructed neurons through the co-registration table (*n* = 15,434 co-registered EM neurons) at CAVE materialization version 1718 [23] and extracted synapses between these cells from the EM reconstruction, excluding autapses. Forty-four fields from 11 of the 12 scans had co-registered neurons, with 4,681 co-registered units from 4,219 EM neurons. We removed pairs of units that map to the same EM neuron and applied no proofreading filter. A directed pair *i → j* was counted as synaptically connected if at least one synapse ran from *i* to *j*. For each area pair or layer pair, EM connection density is the fraction of directed co-registered pairs with at least one synapse. For cell-type pairs (**Figure 3A**), EM synapse density is the number of synapses between two cell types per ordered neuron pair. It uses all 12,694 co-registered neurons of the 15 excitatory types, so it does not depend on which neurons share a field. To compare functional and synaptic connectivity at the single-connection level, we computed the enrichment ratio (risk ratio). This is the synaptic-connection rate among functionally connected pairs (FC+) divided by that among unconnected pairs (FC*−*), computed from per-field counts. Here a pair had a synapse if at least one synapse joined the two neurons in either direction. A pair was FC+ if a state graph had a same-time or lagged edge between the two neurons. We pooled the two state graphs as edge instances, so a pair joined in both counts twice. We resampled the *n* = 44 EM-co-registered fields with replacement (*N* = 5000) to obtain 95% confidence intervals. Confidence intervals and tests of EM density across area pairs and layers also came from a bootstrap over fields, with Holm correction. Comparison edge sets from lagged correlation and Granger causality used the same blocks and the same pair scoring. A lagged-correlation edge joined a pair whose lag-one correlation had *p <* 0.05 (Fisher *z*) in either direction. A Granger-causality edge joined a pair whose lag-one coefficient had *p <* 0.05 in either direction, in one conditional VAR(1) model of all neurons. No method used a multiple-comparison correction. For Figure 1E, we also ran CITS and both comparison methods on the fluorescence traces released with the dataset. These used the same neurons, blocks, states and settings as the deconvolved activity.

Distances are in-plane, because all neurons of a field lie at one imaging depth. We related functional and EM densities across cell-type pairs by Spearman correlation. Confidence intervals came from the field-cluster bootstrap of the functional densities, with EM densities held fixed.

#### Data Aggregation

We summarized each scan’s circuit by two metrics computed on absolute edge weights. Edge density is the fraction of possible directed pairs with a nonzero edge. A same-time edge counts in both directions and a lagged edge in its time order. Edge strength is the mean absolute weight over nonzero edges. We computed these metrics per scan, pooling its fields so that each neuron pair counts once. For a subset, such as an area pair, a layer group or a cell type, we restricted both counts to the subset. The connections of a cell type are the edges between neurons of that type and other typed neurons, in either direction. Although edge sign is interpretable where the dependence is monotone, we aggregated magnitudes because our claims concern the presence and strength of functional connections rather than their polarity.

### 4.6 Statistical Analysis

#### 4.6.1 Data Structure and Unit of Analysis

Each recording session comprised multiple scans, each of which contained eight imaging fields (spatially distinct two-photon fields of view, each capturing a population of neurons at a specific cortical location, referred to as “fields” throughout, following MICrONS dataset conventions). For all state comparisons, the unit of analysis was the scan (Independence of Observations). For each measure and subset, the per-scan change is 100*×*(high*−*low) divided by the low-arousal mean of the subset over scans. We report the mean over the 12 scans with a scan-level 95% *t* interval. We tested it with an exact two-sided sign-flip test over scans, with 2^12^ patterns and a smallest attainable *p* of 0.00049. For comparison, we also tested each state difference without pairing the two states of a scan. This exact two-sample permutation test relabeled the scan values in all possible ways. Its *p* values were corrected within the same families. We also computed a hierarchical bootstrap interval, resampling sessions, scans within sessions and fields within scans (10,000 draws). The number of fields used varied by analysis. *n* = 96 fields contributed to area-level connectivity, *n* = 44 to the cell-type network (fields with EM morphological cell-type labels), all 44 to the EM synapse-enrichment and synaptic-density analyses (fields with EM-reconstructed synapses between co-registered pairs), and *n* = 36 to the layer analysis (fields with sufficient co-registered neurons per cell-type-assigned layer).

##### Independence of Observations

Fields within the same session and scan may share some neurons, potentially violating independence assumptions. They also share blocks and state labels. We therefore used the scan as the unit of every state comparison. The total number of fields analyzed varied by hypothesis, with exact sample sizes reported for each test.

#### 4.6.2 Hypothesis Testing Procedures

##### Area-Specific Decomposition

For each scan, we computed within-area connectivity metrics (edge density, edge strength) separately for each area (V1, LM, AL, RL) and between-area metrics separately for each of the 12 directed between-area pairs. Within-area metrics used submatrices containing only neurons from the same area. Between-area metrics used cross-area submatrices. The primary analysis characterizes within-state organization using the scan-level procedure described above. We computed per-state means over scans for each area pair, and assessed within-state contrasts (within-area vs. between-area) with exact sign-flip tests over scans. We related functional and synaptic areal organization by rank-correlating functional edge density with EM synaptic density across the nine area pairs without LM (Spearman, field-cluster bootstrap test). Each area pair involving LM had at most 865 co-registered neuron pairs and at most one with a synapse, too few for reliable estimates.

##### Within-Area/Between-Area

We classified connections as within-area (same area) or between-area (different areas). We computed mean connectivity per category per scan and arousal state. For paired comparisons (low vs. high arousal within the same scan), and for comparisons of within-area with between-area connections within the same state, we used exact sign-flip tests over scans.

##### Layer-Specific

We assigned neurons to layers by the morphological cell types of the cell-type network below [26], so both analyses share one set of labels. Types L2a to L3b formed layer 2/3, L4a to L4c formed layer 4, and L5ET, L5NP, L5a and L5b formed layer 5. These types exist only for EM-co-registered neurons. Layer 6 was excluded, because only 10 of the 44 EM fields held co-registered layer-6 units (40 in total). We computed mean within-area and between-area connectivity at each layer per scan. For paired comparisons (low vs. high arousal within the same scan and layer), we used exact sign-flip tests over scans. Comparisons between layers within a state used Holm-corrected exact sign-flip tests.

##### Cell-Type Network Graph

To construct the aggregate cell-type functional circuitry network (**Figure 3A**), we assigned neurons to 17 cell types (DTC, Inhibitory, L2a, L2b, L2c, L3a, L3b, L4a, L4b, L4c, L5ET, L5NP, L5a, L5b, L6short-a, L6short-b, L6tall-a) based on co-registration with morphological classification from the EM reconstruction [26]. For each scan, we computed the edge density for each directed cell-type pair, which is defined as the fraction of possible neuron pairs for that cell-type pair with a nonzero inferred CITS connection, pooled over the fields of the scan (*n* = 44 fields with EM co-registration). The network includes excitatory cell types only. Inhibitory neurons were too sparse for cell-type-resolved connectivity, with 35 inhibitory units in total versus over a hundred for most excitatory subtypes, fewer than one per field on average. Cell-type pairs supported by fewer than three co-registered fields were also excluded as small-sample. The mean across scans constitutes the reported edge density for each cell-type pair and arousal state.

##### Predictive Correlation

We computed the Pearson correlation between model predictions and actual responses per neuron and trial. For each neuron we formed a predictive-correlation profile, its per-trial predictive correlation across trials (all trials for edge weight, and the trials of one state for synapse count). Profile similarity between two neurons is the across-trial Pearson correlation of their profiles. For each directed area pair and arousal state, we computed the Spearman correlation between CITS edge weight and profile similarity, and between EM synapse count and profile similarity, over the neuron pairs in that area pair. For edge weight, we computed it within each field, with zero weight for pairs without an edge, and averaged it over fields and then over scans. For synapse count, we computed it over the co-registered pairs of each scan and averaged it over scans. We report means over scans with bootstrap 95% CIs and tested each against zero (exact sign-flip tests over scans, Benjamini-Hochberg-corrected over the tested area pairs of each state, with the same sign required in at least four of five sessions). Synapse-count values from fewer than five scans were not tested. We chose the architecture based on the winning model at the Sensorium 2023 competition [42]. We obtained model predictions using this architecture, which employs an ensemble of 3D convolutional networks with depthwise separable convolutions and temporal attention mechanisms (see Supplementary Methods for architectural and training details). We trained this model architecture on MICrONS data using seven-fold cross-validation (out-of-fold predictions), so that each trial’s prediction comes from a model trained on the other six folds. Evaluation is on these held-out trials only. Model inputs include video frames, pupil center, pupil size, and running speed. The model was not trained to predict arousal state, and fold assignment did not use arousal labels. For Figure 4D, predictive correlation is the Pearson correlation between predicted and recorded responses over the trials of one state. We averaged it over the neurons of each layer or cell type in each scan, and then over scans. Layers in this analysis follow imaging depth. Layer 2/3 spans 150 to 350 *µ*m, layer 4 spans 350 to 450 *µ*m and layer 5 spans 450 to 650 *µ*m. Cell types were the EM cell-type classes of co-registered neurons [69]. We compared groups within each state with the exact sign-flip test over scans. Benjamini-Hochberg correction ran within each state, separately for layers and cell types. A mark also required the same sign in at least four of five sessions.

All connectivity comparisons used non-parametric tests (exact sign-flip tests over scans, fieldcluster bootstrap tests and Spearman correlation) due to non-normal distributions. Predictive correlation related hypothesis tests also used exact sign-flip tests over scans. The EM contrast of within-area and between-area pairs used Fisher’s exact test. All sign-flip tests were two-sided.

##### Robustness rules

A difference could come from one session, from movement or from a few large fields. We therefore called a difference robust only if it met six rules. It needed *q <* 0.05, the same sign in at least four of five sessions, and a stable sign when any one session was left out. After removal of movement and brightness signals, it had to keep its sign and at least half its size. Its sign also had to hold with equal weighting of fields and in both folds. The movement signals were saccade onsets, treadmill speed, eye speed and frame-to-frame brain motion, each filtered with time constants of 0.5, 1 and 2 s. Pupil-center position completed the 14 signals. We regressed them and four imaging-brightness signals from the activity of each neuron within each state and re-estimated the weights. Each fold, one half of the data, had its own graphs and weights.

##### Sensitivity to the controls

Movement differed between the states, so we repeated the full analysis under three designs that control it more strictly. The running-controlled design regressed running history from the pupil index, fitted on stationary frames with a treadmill speed of at most 1 cm/s. Running history was the causal exponential average of treadmill speed with time constants of 1, 3, 10 and 30 s. This design also ended stretches at running, from 1 s before to 2 s after any frame above 1 cm/s, and at treadmill dropouts. The stationary design also removed frames from one frame before to 2 s after each saccade onset. The movement-adjusted design applied the movement and brightness regression above to the stationary design. Each design repeated the selection, the gates, CITS, the weights and the statistics. The stationary design analyzed 11 scans, because its label-validity gate was undetermined in one scan.

##### Further checks

Pooled density gives larger fields more weight, so a field-size check compared smaller and larger fields. It ranked the eight fields of each scan by the number of modeled neurons. It compared the mean per-field density change of the four smaller and the four larger fields. The per-field change was 100 *×* (high*/*low *−* 1) of the field’s density. We computed it for all edges, for same-time edges and for lagged-only edges, meaning ordered pairs joined only across frames. We tested the per-scan difference with the exact sign-flip test over 12 scans. Because activity was lower in high-arousal periods, the rate control restricted the analysis to neurons whose rate changed by less than 10%. The eye-movement analysis compared faster and slower eye movement within each state. It split blocks at the median eye speed within pupil-index quintiles. To test whether the state result is specific to CITS, we repeated it with other measures. The mean absolute correlation over all neuron pairs took the larger of the zero-lag and lag-one values. We also computed it over correlation edges, and standardized pairwise and conditional Granger-causality weights over their own edges. Edges of each method had *p <* 0.05 in its own test.

##### Multiple Comparisons Correction

We performed multiple statistical tests across different hypotheses. To control the false discovery rate (FDR), we applied the Benjamini-Hochberg procedure [70] within each hypothesis family. Families were defined as follows. The six network rows formed one family. These were density and edge weight for all, within-area and between-area pairs. The 16 ordered area pairs, the six layer groups and the cell-type groups formed one family per measure. Predictive similarity formed one family per state over its tested area pairs. This family-wise approach is appropriate because each family addresses a distinct scientific question. Comparisons among areas or among layers within a state used Holm correction. We classified network rows as primary, area pairs and layer groups as secondary, and cell types as exploratory. For all hypotheses with multiple tests, we report FDR-corrected *p*-values in the main text. For single-test hypotheses, no correction is needed and we report uncorrected *p*-values. Tests with FDR-corrected *p <* 0.05 (or *p <* 0.05 for single tests) were considered significant. We pre-specified an analysis of fluorescence-based edge weights before computing any state contrast. Its result was inconclusive, and all analyses reported here were specified afterwards. We chose the minimal-control design as the main analysis before computing it, and report the other designs as sensitivity analyses.

##### Effect Sizes

For correlation analyses, we report Spearman or Pearson correlation coefficients as appropriate. For differences between conditions, we report percent changes or raw differences (Δ) along with the relevant test statistic and *p*-value (exact sign-flip tests over scans).

We implemented the analyses in Python.

## Contributors

RB and RA-A designed the study and conceptualized the analysis framework. RB performed the analysis with contributions from RA-A. RB, JZ, YW, and HWS curated the data and implemented the software. RB and RA-A interpreted the findings with contributions from SM. RA-A supervised the study and acquired funding. RB wrote the first draft with contributions from RA-A. All authors reviewed and approved the manuscript.

## Declaration of interests

The authors declare no competing interests.

## Acknowledgments

We thank members of the Abbasi Lab for their insightful feedback on this work. This study was partially supported by the National Institute of Mental Health of the National Institutes of Health under award number RF1MH128672 (model development and initial validation) and by the National Library of Medicine of the National Institutes of Health under award number R01LM014619 (cell-type and layer-specific analysis). The content is solely the responsibility of the authors and does not necessarily represent the official views of the National Institutes of Health. During this work the authors used language models as an aid in writing analysis code, searching the literature, and editing the manuscript text. The authors directed this use, reviewed and verified all code, results, references, and text, and take full responsibility for the content of the publication.

## Data and Code Availability

Analysis code will be made publicly available at https://github.com/abbasilab/mfc upon publication. Raw calcium imaging data and electron microscopy reconstructions are available through the MICrONS project (https://www.microns-explorer.org/). CITS is described in [27] and is available at https://github.com/abbasilab/cits. It builds on the time-aware PC framework [9], available at https://github.com/shlizee/TimeAwarePC.

## Supplementary Methods

### Model Architecture and Training

For predictive correlation analysis, we chose the architecture based on the winning model at the Sensorium 2023 NeurIPS competition [42, 71], which achieved first place in predicting large-scale mouse visual cortex activity from videos. The model employs an ensemble approach combining multiple architectural innovations.

### Architecture Components

The base architecture consists of 3D convolutional networks with depthwise separable convolutions [72], enabling efficient spatiotemporal feature extraction from video inputs. The model incorporates temporal attention mechanisms [73] to capture long-range dependencies across time. The ensemble combines predictions from multiple models trained with different configurations, improving robustness and generalization.

### Training on MICrONS Data

We trained this model architecture on the MICrONS dataset, using the same visual stimuli (natural movies and synthetic parametric videos) and behavioral signals (pupil center, pupil size, running speed) that were available during calcium imaging. We divided trials into seven folds for out-of-fold predictions, so that each trial’s prediction comes from a model trained on the other six folds. This ensures that all predictions were evaluated on held-out data. We trained the model to predict neural responses (calcium fluorescence traces from the MICrONS processed dataset) from video frames and behavioral inputs (pupil center, pupil size, running speed). The model learns a mapping from these inputs to neural activity patterns.

### Relationship to Arousal

Model inputs include video frames, pupil center, pupil size, and running speed (i.e., the model can use moment-to-moment state when predicting responses). We performed the train/test and fold splits at the trial level without regard to arousal state. We did not train the model to predict arousal state. We applied trial-level arousal-state labels only after obtaining predictions.

### Performance

On the original Sensorium 2023 competition dataset, this architecture achieved single-trial correlations of 0.2913 on the main track and 0.2215 on the bonus track (final test set) [71]. On the MICrONS data used in this study, we computed the same single-trial correlation metric as in the Sensorium competition (without regard to arousal state). The metric is the mean correlation between predicted and actual responses per trial, averaged over trials and units. The model achieved a single-trial correlation of 0.2878 on out-of-fold predictions and 0.3198 on the held-out test set (752 trials). The model’s ability to capture stimulus-response relationships makes it well-suited for assessing predictive correlation as a measure of how well neural responses can be predicted from the visual input.

### Predictive correlation: full pipeline

For transparency and reproducibility, the end-to-end pipeline is as follows.

1. *Data.* MICrONS calcium imaging data: for each session and scan, we use processed calcium fluorescence traces (responses) and the corresponding inputs for every trial. Inputs to the model are video frames, pupil center, pupil size, and running speed. Target is the trial-wise response matrix (units *×* time). The unit of analysis is session-scan-unit (146,388 units across 16 session-scans).
2. *Train/test split and out-of-fold predictions.* We held out a subset of 752 trials from fold assignment and did not use them in training (separate test set). We split the remaining trials (6,672 across 16 session-scans) into *K* = 7 folds. For each fold, the model is trained on the other six folds (*∼*86% of those trials) and predicts on the held-out fold (*∼*14%). Thus each fold trial receives a prediction from a model that did not see that trial during training (out-of-fold predictions). The split is at the trial level, is identical for all neurons within a session-scan, and does not use arousal state or any trial metadata. Reported predictive correlations are computed from out-of-fold predictions.
3. *Model and training.* We chose the model architecture based on the winning model at the Sensorium 2023 competition. It is a depthwise separable 3D CNN (“dwiseneuro”) with temporal attention and approximately 309 million trainable parameters, trained to predict neural responses from video and behavioral inputs (pupil center, pupil size, running speed). Training minimizes Poisson negative log-likelihood (NLL) between predicted and actual responses on the training trials of each fold. The optimizer is AdamW with base learning rate 3 *×* 10*^−^*^4^ (with warmup and cosine decay to 1% of base), weight decay 0.05, batch size 32. Training runs for 21 epochs (3-epoch linear warmup followed by 18 epochs with cosine learning-rate decay). The held-out fold is used only for monitoring validation loss and validation correlation and is never used for training or early stopping. We performed training on two NVIDIA RTX 5000 Ada Generation GPUs. Each fold took approximately 12 hours. Model inputs include video, pupil center, pupil size, and running speed. Arousal-state labels are not used for fold assignment, as a training target, or in model selection.
4. *Evaluation.* For each trial, the trained model (or ensemble) produces a predicted response matrix of shape (units *×* time). Predictions are saved per trial so that each trial has a matching pair (predicted trace, actual trace) for every unit. The out-of-fold predictions are used in the arousal analysis reported in the main text.
5. *Arousal-state assignment and predictive correlation.* After all predictions are obtained, each trial is labeled low-arousal or high-arousal as described in Methods. For each scan and each unit, we collect all low-arousal trials and all high-arousal trials. Within each state, we concatenate the predicted and actual traces across trials, yielding one predicted and one actual trace per unit per state. The predictive correlation in that state is the Pearson correlation between these concatenated traces. Arousal-state labels were not used for fold assignment or as a training target.

### Training dynamics and convergence

We ran training for 21 epochs with no early stopping across seven cross-validation folds. Loss decreased and validation correlation increased over the training epochs. The results reported here use out-of-fold predictions for the 12 analyzed scans.

## Supplementary Tables

Supplementary Table S1. gives the statistics behind the marks in Figures 2D, 3C and 3D. Supplementary Table S2 gives the edge-weight change under stricter designs and with other methods.

**Table S1:** Statistics of the between-state changes shown in the main figures. Change from low to high arousal in edge weight, or edge density where stated, in the main analysis. The 95% CI is the *t* interval over scans. In the layer-4 between-area row, one scan with a 44% increase widens the interval to include zero. Scans gives how many scans changed in the direction of the mean. *q* is the Benjamini-Hochberg value within each family. Every change with *q <* 0.05 also meets the other five robustness rules in Methods, except edge density over all pairs and within areas. Their sign reverses when fields are weighted equally. Cell types are exploratory. DTC was tested but is not shown in Figure 3D.

| Figure | Group | Change | 95% CI | Scans | *q* |
|---|---|---|---|---|---|
| Network | Edge density, all pairs | −2.4% | −3.3% to −1.4% | 11 of 12 lower | 0.001 |
| Network | Edge density, within area | −1.6% | −2.7% to −0.6% | 10 of 12 lower | 0.008 |
| Network | Edge density, between areas | −4.1% | −6.0% to −2.1% | 11 of 12 lower | 0.003 |
| Network | Edge weight, all pairs | +9.4% | +7.1% to +11.7% | 12 of 12 higher | 0.00098 |
| Network | Edge weight, within area | +9.8% | +7.4% to +12.3% | 12 of 12 higher | 0.00098 |
| Network | Edge weight, between areas | +8.2% | +6.1% to +10.3% | 12 of 12 higher | 0.00098 |
| 2D | AL→AL | +9.6% | +5.5% to +13.7% | 11 of 12 higher | 0.003 |
| 2D | AL→LM | +2.8% | −5.5% to +11.0% | 5 of 9 higher | 0.46 |
| 2D | AL→RL | +7.6% | +4.4% to +10.8% | 11 of 12 higher | 0.002 |
| 2D | AL→V1 | +9.2% | +5.6% to +12.9% | 12 of 12 higher | 0.001 |
| 2D | LM→AL | −4.5% | −12.1% to +3.0% | 5 of 9 lower | 0.24 |
| 2D | LM→LM | +4.5% | +1.6% to +7.3% | 11 of 12 higher | 0.012 |
| 2D | LM→RL | +4.7% | −0.7% to +10.0% | 8 of 12 higher | 0.11 |
| 2D | LM→V1 | +8.1% | +6.1% to +10.2% | 12 of 12 higher | 0.001 |
| 2D | RL→AL | +7.1% | +4.3% to +9.9% | 11 of 12 higher | 0.002 |
| 2D | RL→LM | +2.3% | −3.1% to +7.7% | 8 of 12 higher | 0.40 |
| 2D | RL→RL | +5.9% | +3.7% to +8.1% | 11 of 12 higher | 0.002 |
| 2D | RL→V1 | +9.0% | +6.3% to +11.8% | 12 of 12 higher | 0.001 |
| 2D | V1→AL | +10.4% | +7.0% to +13.9% | 12 of 12 higher | 0.001 |
| 2D | V1→LM | +8.3% | +6.1% to +10.5% | 12 of 12 higher | 0.001 |
| 2D | V1→RL | +8.2% | +5.5% to +10.9% | 12 of 12 higher | 0.001 |
| 2D | V1→V1 | +10.6% | +8.0% to +13.2% | 12 of 12 higher | 0.001 |
| 3C | L2/3 within area | +7.5% | +3.5% to +11.4% | 10 of 11 higher | 0.012 |
| 3C | L4 within area | +7.6% | +2.4% to +12.7% | 10 of 11 higher | 0.023 |
| 3C | L5 within area | +9.5% | +2.4% to +16.7% | 8 of 10 higher | 0.026 |
| 3C | L2/3 between areas | +11.9% | +6.6% to +17.2% | 10 of 11 higher | 0.012 |
| 3C | L4 between areas | +9.3% | −1.6% to +20.2% | 8 of 9 higher | 0.023 |
| 3C | L5 between areas | +7.2% | −2.2% to +16.5% | 5 of 7 higher | 0.11 |
| 3D | DTC | +21.6% | +12.6% to +30.6% | 4 of 4 higher | 0.17 |
| 3D | L2a | +5.0% | −5.3% to +15.2% | 6 of 8 higher | 0.35 |
| 3D | L2b | +3.9% | −10.5% to +18.2% | 6 of 7 higher | 0.55 |
| 3D | L2c | +8.5% | −1.3% to +18.2% | 7 of 9 higher | 0.15 |
| 3D | L3a | +9.1% | +5.9% to +12.3% | 10 of 11 higher | 0.012 |
| 3D | L3b | +10.1% | +4.1% to +16.1% | 9 of 10 higher | 0.023 |
| 3D | L4a | +8.2% | +3.3% to +13.0% | 9 of 10 higher | 0.023 |
| 3D | L4b | +9.9% | +5.7% to +14.1% | 10 of 11 higher | 0.012 |
| 3D | L4c | +4.1% | −4.8% to +13.0% | 7 of 10 higher | 0.35 |
| 3D | L5ET | +9.1% | +2.1% to +16.2% | 7 of 9 higher | 0.056 |
| 3D | L5a | +6.8% | +0.9% to +12.7% | 8 of 9 higher | 0.070 |
| 3D | L5b | +8.3% | −0.6% to +17.2% | 7 of 8 higher | 0.12 |

**Table S2:**
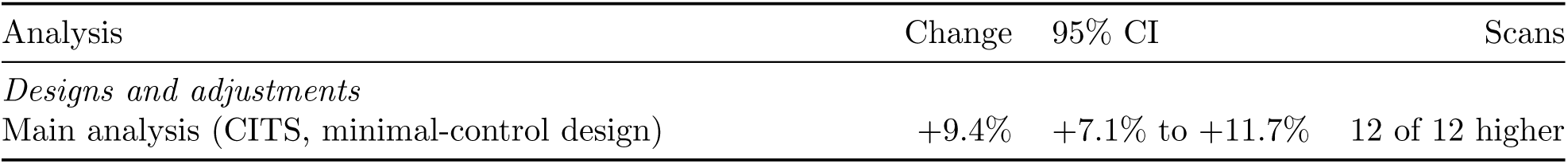
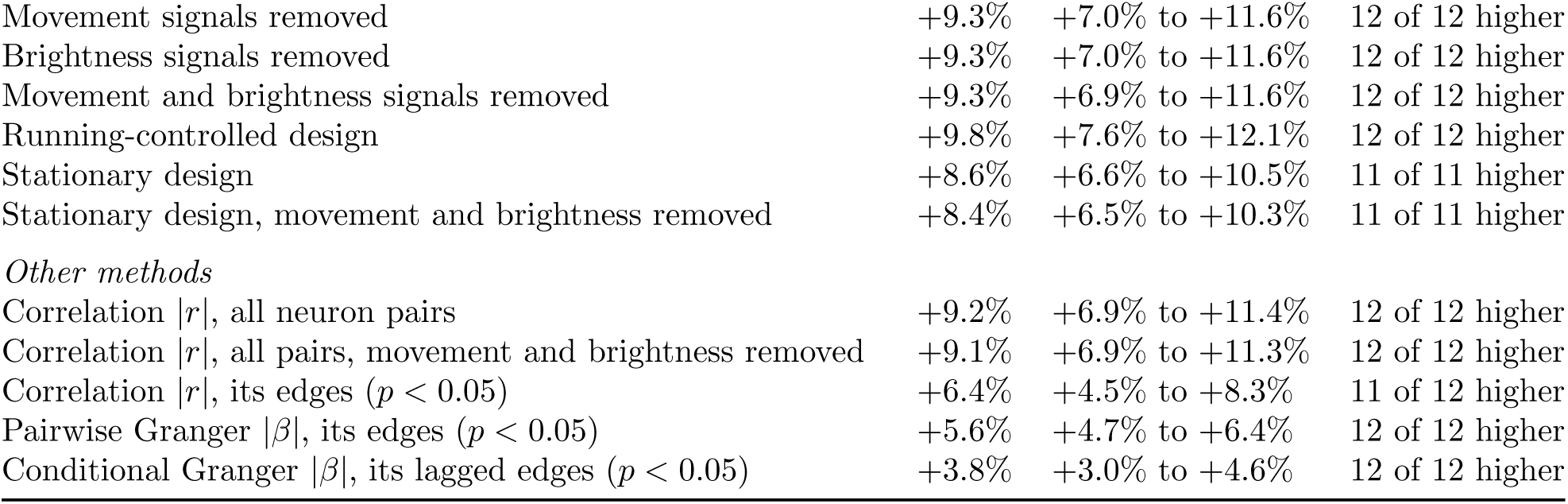
Edge-weight change under stricter designs and with other methods. Change from low to high arousal over all neuron pairs. Designs and adjustments are defined in Methods. Other methods use the same matched blocks. Area, layer and cell-type comparisons used CITS only. The 95% CI is the *t* interval over scans. The stationary design has 11 scans.

## Supplementary Figures

**Figure S1:**
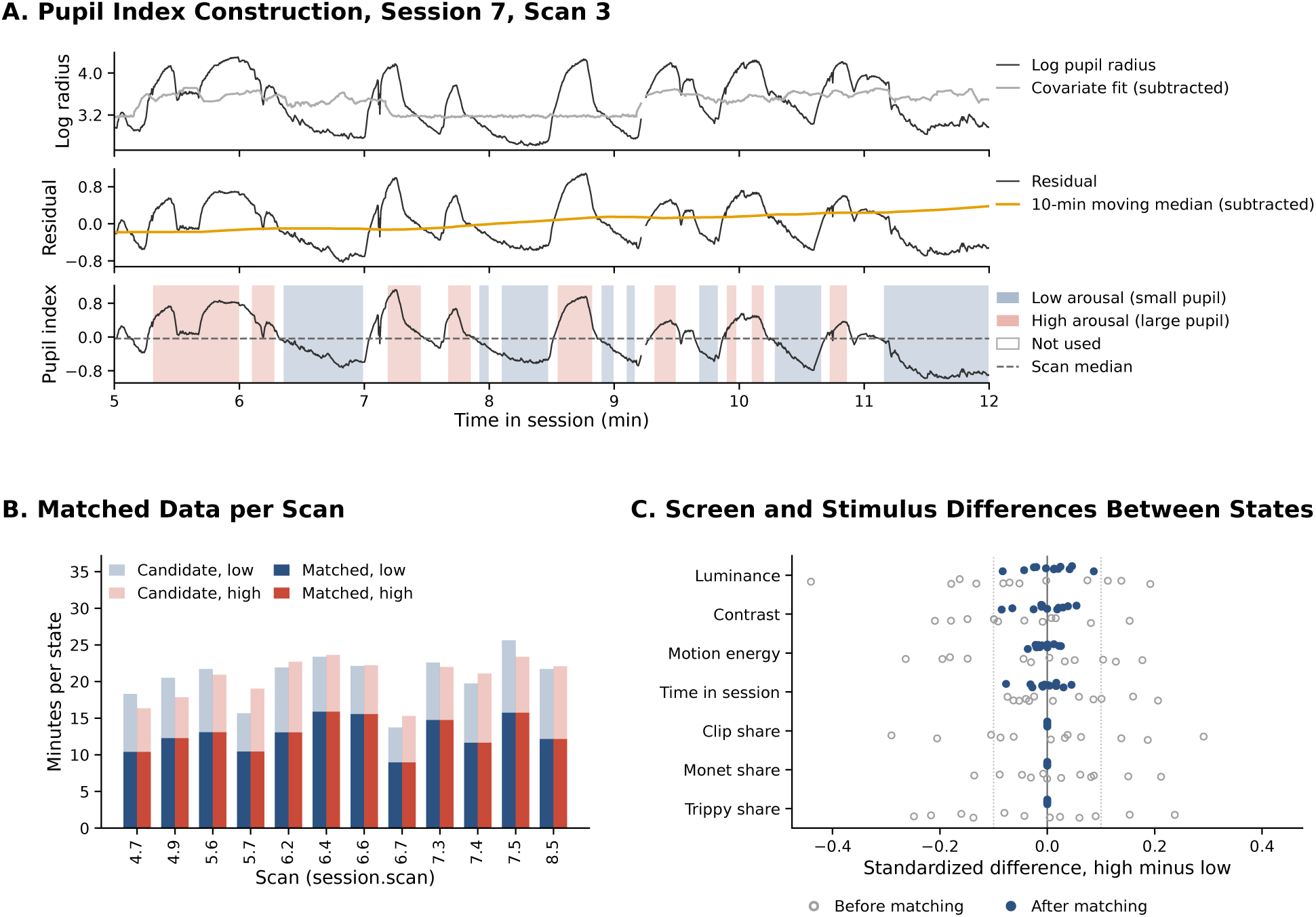
Arousal-state definition and matching. **(A)** Construction of the pupil index in one scan (session 7, scan 3). Top, log pupil major radius and its fit by stimulus covariates. Middle, the residual and its 10-minute moving median. Bottom, the pupil index, the residual minus its moving median, with higharousal and low-arousal periods. Unshaded stretches are not used. **(B)** Candidate and matched blocks per scan and state. Matched blocks are equal in both states (14,546 per state, 154 minutes). **(C)** Standardized differences in screen covariates and stimulus shares between states, before and after matching. Dotted lines mark *±*0.1. After matching, per-scan differences are at most 0.09 and stimulus shares are identical.

## Notes

### Competing Interest Statement

The authors have declared no competing interest.

### Summary of Updates

We revised the analysis, figures and text. We now define arousal states by a pupil index within each scan and match them for visual stimulus and session time. We now infer connections from deconvolved activity rather than fluorescence. The neuron count now includes only the 12 analyzed scans. Statistical tests now treat each scan as one sample. The electron microscopy comparison now checks consistency with synaptic anatomy and no longer ranks inference methods. We removed the claims about layer 6 and spatial extent, which the revised analysis did not support.

